# Germ-stem cells and oocyte production in the Honeybee Queen Ovary

**DOI:** 10.1101/2022.09.22.509099

**Authors:** Georgia Cullen, Joshua B. Gilligan, Joseph G. Guhlin, Peter K. Dearden

**Affiliations:** Laboratory for Evolution and Development, Biochemistry Department, University of Otago, Dunedin, Aotearoa-New Zealand; Biological Heritage National Science Challenge, Biochemistry Department, University of Otago, Dunedin, Aotearoa-New Zealand; Genomics Aotearoa, Biochemistry Department, University of Otago, Dunedin, Aotearoa-New Zealand

**Keywords:** Honeybees, Ovary, Cell Division, Reproductive rate, Gene expression

## Abstract

Understanding the reproduction of honeybee queens is crucial to support populations of this economically important insect. Here we examine the structure of the honeybee ovary to determine the nature of the germ-stem cells in the ovary. Using a panel of marker genes that mark somatic or germ-line tissue in other insects we determine which cells in the honeybee ovary are somatic and which germline. We examine patterns of cell division, and demonstrate that, unlike Drosophila, there are no single germ-line stem cells that provide the germ-line in honeybees. Germ-line stem cells are clustered in groups of 8 cells, joined by a polyfusome, and collections of these, in each ovariole, maintain the germ-line during reproduction. We also show that these 8-cell clusters can divide, and that their division occurs such that the numbers of germ-line stem cells are relatively constant over the reproductive life of queen honeybees. This information helps us to understand the diversity of structures in insects reproduction, and provide information to better support honeybee reproduction.

## Introduction

Honeybees are economically important insects, providing hive products such as honey, wax and propolis, as well as pollination services estimated at US$34 billion/ year in 2012^1^. Because of their eusocial life history strategy^2^, honeybee reproduction depends, to a large extent, on a single queen bee present in each hive. Reproduction, and thus productivity of a bee colony depends on the activity of the queen ovary. Given recent challenges to both managed and unmanaged honeybee populations^3^, such as varroa mite vectored disease and the widespread use of insecticides, it is important to know how best to support honeybee queen reproduction.

Despite the importance of honeybees, little is known about the structure and function of honeybee ovaries. We do not understand the control of stem cell division in the ovary, or even where the germstem cells are placed. Knowing more about the structure and function of the queen ovary may help us determine the best time to intervene in honeybee queen development to maximise the rate of reproduction, quality of offspring, and total reproductive output.

Insect ovaries are categorised into structural groups^4^. Honeybees have polytrophic meroistic ovaries^5^, meaning the oocyte is nourished by a set of nurse cells (trophocytes)^4^. These are attached to the oocyte via cytoplasmic bridges (polyfusomes and ring canals^6^) that travel with each oocyte down the ovariole before dying just before egg laying.

Honeybees are holometabolous insects and their polytrophic meroistic ovary structure have meant they are often compared to *Drosophila melanogaster* which has a superficially similar ovary structure^7^. It is clear that even amongst this ovary type and holometabolous insects themselves, the structure and function of the ovary differ. This is no surprise given the different life-history contexts these species exist in; honeybee queens, in particular, are reputed to be able to lay between 500 and 3000^8–10^ eggs a day and to live 1-5 years^11,12^ while *D. melanogaster*, in a laboratory environment, lay an average of 615 offspring^13^ over their approx. 90-day life^14^.

In honeybees, part of this reproductive output is explained by the number of ovarioles each queen possesses. Ovarioles are the functional units of the ovary, each being a long strand of cells. At the anterior end is the terminal filament, and then, towards the posterior, the germarium, followed by the vitellarium, from which mature eggs are released. Experiments to determine the range and number of ovarioles in honeybee ovaries indicate they contain between 233 and 438 (median 320) in each of two ovaries^15^, whereas in *Drosophila*, the number of ovarioles varies between 18 and 20, in *Tribolium* (red flour beetle) it is 8, in *Bombyx mori* (silk moth) it is 8, and in non-Apis Hymenoptera, numbers range from 2-10,000 ovarioles, with queen ants and bees making up the larger numbers, but with most species having between 8 and 100^16^.

In *Drosophila*, the germarium in each ovariole contains two germ stem cells whose division is crucial to reproductive output; reviewed in^17^. These single cells divide with one daughter cell remaining a stem cell, and the other forming the first stage of oocyte production, a cystocyte. As cystocytes divide, they remain attached by a cytoplasmic bridge, named a polyfusome, which links these cells in a cyst. One of the cells of a cystocyte undergoes meiosis I, leaving a presumptive oocyte attached to a cluster of cells, now named nurse cells or trophoblasts, via a series of cytoplasmic links named ring canals. As the oocyte develops, these nurse cells divide and endoreplicate their genomes, becoming polyploid, and generate proteins and RNA which are transported into the oocyte, both providing nutrition and patterning the oocyte. As the oocyte matures and is readied to be laid, the nurse cells contract, dumping their cytoplasm into the oocyte, and dying; this provisioned and patterned egg is then laid.

While this process has been well studied in *Drosophila*, the ovaries of *Apis* appear to have significant differences from the *Drosophila* model. Previous studies in honeybees have suggested that germ stem cells are present in the terminal filament of the ovary^5^, as has been described for another bee species Osmia^18^. Histochemical and contrast studies indicate that the germarium of honeybees is filled with resting germ cells, but it is not clear how these cells are arranged and whether they divide^4^. At the boundary of the germarium and the vitellarium, it is reported that the cells are arranged into rosettes, joined by a polyfusome, prefiguring the development of a nurse/oocyte cluster^19,20^. The cysts in *Apis* also form differently from *Drosophila*, with each nurse cell cluster separated by a narrow neck from the oocyte^19,21,22^.

To begin to understand the placement of germ-line and somatic cells in the honeybee ovary, and the source of its remarkable fecundity, we examined the ultrastructure of the ovariole. We also classified cell types in the ovary as somatic or germ-line and examined the placement and timing of cell divisions in the ovary as a way to understand reproduction in honeybees.

## Materials and Methods

### Honeybees

Actively laying *Apis mellifera* queens were selected from different hives across Dunedin, Otago. Replicates of each experiment were carried out with queens that were not closely related. For immunohistochemistry experiments, tissues came from 5 replicate queens. EdU experiments used 5 replicates for 24-hour experiments and 3 for 48-hour experiments. *In situ* hybridisation involved 18 replicate queens, at least 3 per gene. Multiple ovarioles for each queen were examined in each case. For polyfusome counting experiments, six queens were divided into two groups (three per group), based on their age. Old queens are defined as being in their second season, having over-wintered and being older than one year, and young queens as in their first season, 3-10 months old. Multiple ovarioles from each queen were examined

### Dissection and Fixation

Laying adult queen honeybees were dissected as soon as possible after hive removal. Unlike previous experiments^21^, and due to concerns about tissue damage, neither bees nor ovaries were ever subjected to freezing unless fixed and stored in methanol. Queens were anaesthetised by cold until movement ceased and then the head was removed from the thorax. The ovaries were dissected into 1x PBS (Phosphate buffered Saline) under a dissecting microscope. As the ovarioles were separated and the intima (eternal membrane on each ovariole) peeled off with fine forceps, they were placed in a microcentrifuge tube with 1x PBS on ice. Once the ovarioles were separated and peeled, the samples were fixed in 1:1 heptane:4% formaldehyde in 1x PBS for 15 minutes, and then rinsed in 0.1% PTx (PBS + 0.1% Triton X-100) 3 times. Ovaries were either used immediately or stored in 100% methanol in the dark at -20°C.

### Identification of honeybee marker genes

Candidate genes were selected based on their developmental functions in mainly *Drosophila*, but also previous studies in *Apis* and other Hymenopterans. See Table 1 for further details. Honeybee orthologues were initially identified by reciprocal best Blastp^23^ hit and then further analysed using Bayesian phylogeny techniques, using MrBayes 3.2.7a^24^. Various models of amino acid evolution were chosen after initial experiments using mixed models. Castor, dpp, tapas and hh were analysed with a WAG model^25^, Traffic jam, Mad6, unzipped and eyes absent with a Jones model^26^, ova with a Dayhoff model^27^ and Bark beetle with a Blosum model^28^.

**Table 1:** Potential markers for somatic vs germ-line fate tested in this study.

| Gene name | Predicted Germline or Somatic? | Expression in *Drosophila* ovary | Expression in other insect ovaries | references |
| --- | --- | --- | --- | --- |
| *vasa* | germline | germline | germline (all holo- and hemimetabolous insects examined) | For examples see <sup>22,36–47</sup> |
| *nanos* | germline | germline | germline (all holo- and hemimetabolous insects examined) | For examples see <sup>38,39,41,48–58</sup> |
| *Bark-beetle* | germline | absent | germline (*Nasonia*) | *Drosophila*<sup>59</sup><br>*Nasonia*<sup>60</sup> |
| *ovo* | germline | germline | not clear in *Bombyx* | *Drosophila*<sup>61</sup>,<br>*Bombyx*<sup>62</sup> |
| *MAD6 (daughters against dpp)* | germline | germline stem cells | no data | *Drosophila*<sup>63</sup> |
| *Traffic-jam* | somatic | somatic cells in contact with the germline | no data | *Drosophila*<sup>64</sup> |
| *Castor* | somatic | follicle cells | no data | *Drosophila*<sup>65,66</sup> |
| *Eyes-absent* | somatic | follicle cells | follicle cells (*Tribolium*) | *Drosophila*<sup>65–67</sup><br>*Tribolium*<sup>68</sup> |
| *Decapentaplegic* | somatic | follicle cells | follicle cells (*Ceratitis capitata*) | *Drosophila*<sup>63,69</sup><br>*Ceratitis*<sup>69</sup> |
| *hedgehog* | somatic | terminal filament and cap cells | no data | *Drosophila*<sup>70</sup> |
| *Unzipped* | somatic | Described as somatic cells of the ovary | No data | *Drosophila*<sup>59</sup> |

### Hybridisation Chain Reaction (HCR)

Stored ovaries were rehydrated over a methanol series: 75%, 50%, 25% methanol/PTw (PBS + 0.1% Tween 20), and then rinsed three times in PTw for 5 minutes at each step. Rehydrated, or freshly fixed ovarioles were permeabilized in PTx for 2 hours at room temperature on a nutator.

Samples were pre-hybridized in 500 µL of 30% probe hybridisation buffer (2.4 M Urea, 5x sodium chloride sodium citrate (SSC), 9 mM citric acid (pH 6.0), 0.1% Tween 20, 50 µg/mL heparin, 1X Denhardt’s solution, 10% dextran sulphate), for 30 minutes at 37°C. Four hundred µL of the hybridisation buffer was removed and the probes were added to the remaining 100 µL of hybridisation buffer and samples at 37°C in one of the following concentrations: 2 µL of 1 µM probe (*Nanos, Castor)*, 2 µL of 2 µM (odd and even) probe (*Vasa)*, 4 µL of 1 µM probe (*Tapas, Bb, TJ, Ovo, Eya)*, 10 µL of 1 µM probe (*MAD6, DPP, Hh)*, or 10 µL of 2 µM probe (*Unzipped)* and the sample was incubated overnight (14-18 hours) to five days at 37°C. The probes were washed 4x for 15 minutes each with 200µL of probe wash buffer (2.4 M Urea, 5X SSC, 9 mM citric acid (pH 6.0), 0.1% Tween, 50 µg/mL heparin) at 37°C and then washed 3x for 5 minutes each with 500 µL of 5x SSCT (5X SSC, 0.1% Tween 20) at room temperature.

Samples were incubated in 500 µL of amplification buffer (5X SSC, 0.1% Tween 20, 10% dextran sulphate) for 30 minutes at room temperature, while the hairpins were prepared by snap cooling 2 µL, of a 3 µM stock (6 pmol in 100 µL of buffer), per hairpin (keeping h1 and h2 separate) – heating to 90°C for 90 seconds, and cooling to room temperature in the dark for 30 minutes. Amplification buffer was removed and the snap-cooled hairpins (h1 and h2) were added in 100µL of amplification buffer at room temperature. The pre-amplification buffer was replaced with the hairpin mixture, and the samples were incubated overnight (14-18 hours) at room temperature in the dark.

The hairpins were removed by washing the samples with 500 µL of 5x SSCT at room temperature for 5 minutes x2, 30 minutes with 0.5 µL of DAPI [4’, 6-Diamidino-2-Phenylindole, Dihydrochloride [D1306]] per 1ml PTw, 30 minutes without DAPI, and five minutes with gentle rocking. The sample was rinsed 3x PTw and then stored in 70% ultrapure glycerol at 4°C in the dark.

### Immunohistochemistry

Ovarioles were used immediately after fixation for immunohistochemistry.

The fixed ovarioles were left in PTx for 2 hours on a rocker for permeabilisation. Then they were put in 500 µL blocking solution (PBS +0.1% Triton X-100 + 5% Normal goat serum + 0.2% BSA) for 30 minutes on the rocker at room temperature, or overnight in the fridge. The ovarioles were then put in primary antibody solution ((1:200 α-mouse-ph3 (Anti-Histone H3 (phospho S10) antibody [abcam 14955]): PBTB) overnight in the dark in the fridge.

The antibody solution was replaced with PTx and washed 4x on the rocker for 15 minutes and the ovarioles were re-blocked in blocking solution (PBTB) for 30 minutes. The blocking solution was replaced with secondary antibody solution (1:1000 goat-α-mouse 488:PBTB) and left in the fridge overnight. The antibody solution was replaced with PTx and washed 4x on the rocker in the dark for 15 minutes. The tissue was then counterstained with phalloidin [Invitrogen Alexa Fluor488 phalloidin [A12379], Invitrogen Alexa Fluor555 phalloidin [A34055]] (3 µL of a 66 µM solution per 100 µL PTx) for 15 minutes on a rocker. The samples were rinsed with PTx and then counterstained for DAPI (0.5µL per 1ml of PTx) for ten minutes. The tissue was finally rinsed 3x with PTx and then stored 70% glycerol in the fridge.

### EdU staining

Queen bees were removed from the hive and carefully pushed, abdomen first, using a cotton wool pad, into a microcentrifuge tube modified with holes made in the bottom. A Hamilton 50 µL syringe fitted with a steel 22s needle with a bevelled tip was inserted shallowly between the tergites of the 4th and 5th segment dorsally and used to inject 8-10 µL of 10µM EdU (16-20 pmol), or until the abdomen visibly expanded. The queen was then put in a queen cage and returned to the hive to resume laying for either 24 hours or 48 hours. After the laying period, the queen was euthanized, and the ovaries dissected, separated and fixed as described above. Fixed ovarioles were used immediately, undergoing permeabilisation in PTx for an hour on the rocker at room temperature. The tissue was blocked for at least 30 minutes at room temperature in PBTB.

Tissue was treated and stained using the instructions of the Click-iT EdU cell proliferation kit (Thermofisher). The tissue was counterstained with DAPI (0.5 µL DAPI (10 mg/ml) per 1ml PTx) for 15 minutes, before being rinsed 3x with PTx. Lastly, the samples were stored in 70% glycerol in the dark at 4°C.

### Microscopy

Confocal was performed using an Olympus FV3000 confocal microscope. A combination of z stacks of up to 80 slices and single slice images were taken.

### Polyfusome counting and statistics

Ovaries were used fresh and stained with phalloidin and DAPI as described above. Images were taken using confocal microscopy on an Olympus FV3000, of the early germarium encompassing each set of polyfusomes. Polyfusomes were counted blind by three staff experienced in insect ovary microscopy.

Means were compared between the older queen group and the younger queen group using an unpaired t.test in python using SciPy 1.7.0 stats.ttest_ind function, omitting NaN^29^.

## Results

### Ultrastructure of the queen honeybee ovariole

Confocal microscopy of DAPI and Phalloidin stained honeybee queen ovarioles (Figure 1) is consistent with previous studies^21,22^ and the morphology of other insect ovarioles. At the anterior end of the ovariole is the terminal filament, made up of initially a single, and then a double, stack of cells with flattened nuclei (Figure 1). These cells appear to separate into a sheath of follicle cells around the germarium. The germarium itself is made up of groups of 8 cells joined by a polyfusome, which stains strongly with phalloidin (Figure 1). The posterior of the vitellarium is marked by the polyfusomes being replaced with ring canals, also strongly staining with phalloidin. At this point, in some cells, chromosomes can be seen, suggesting cell divisions of some sort are occurring in this region. We propose this is perhaps where meiosis I is beginning (Figure 1), but staining with markers of meiotic cells will be required to confirm this. Oocytes remain in Metaphase I of meiosis until egg laying^30^ Posterior to this boundary, in the vitellarium, presumptive oocytes become larger than surrounding cells. These come to be surrounded by follicle cells and move into separate chambers from their attached nurse cells, which become larger, and more numerous over this time (as in other hymenoptera^31^) Finally the oocyte and nurse cells, still attached, are in separate chambers, the oocyte surrounded by follicle cells and filling with yolk granules. The oocyte continues to grow until the nurse cells collapse and dump their cytoplasm into the oocyte, releasing it for laying. The entire ovariole is surrounded by a tough membrane, the intima, which must be removed in most cases to improve staining, and, through its structure, appears to be a contractile tissue.

**Figure 1:**
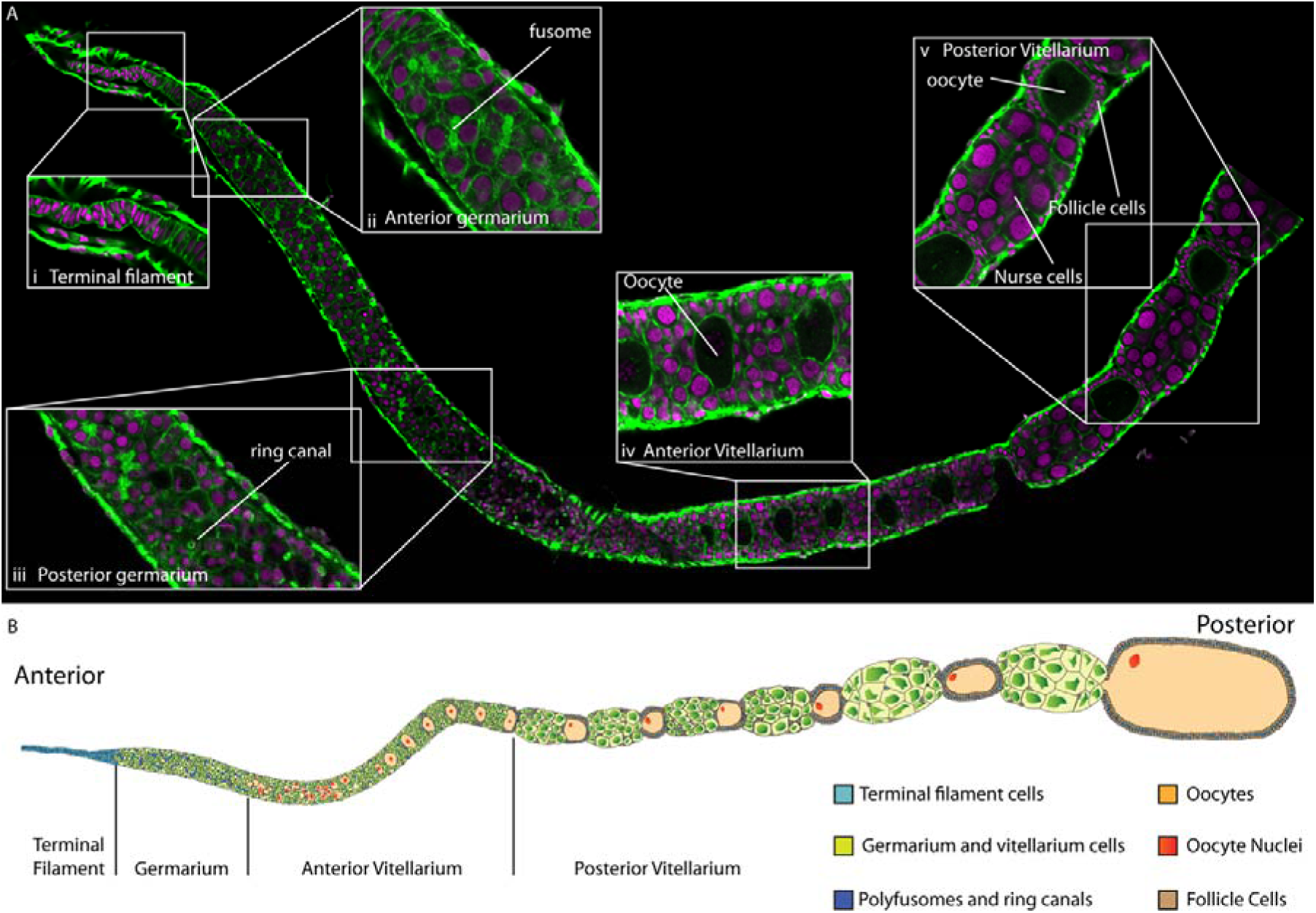
Ultrastructure of the honeybee queen ovariole. A) Composite confocal micrograph of a single honeybee queen ovariole stained with DAPI (purple) and Phalloidin (Green). Insets show key regions of the ovariole. i) Terminal filament showing flattened cells and nuclei ii) Anterior germarium, showing the transition from the flattened cells of the terminal filament to rounded cells and nuclei and polyfusomes staining with phalloidin. iii) Posterior germarium, here polyfusomes change into ring canals (circular phalloidin stained structures, in the same region we see DAPI stained chromosomes in a small patch of cells indicating meiosis or mitosis is occurring. iv) Anterior vitellarium. Here clear oocytes (marked by large size and very faint, or absent nuclear staining), line up in the ovariole. v) Posterior vitellarium, the oocyte and nurse cell group are separated, and follicle cells form a columnar epithelium around the oocyte. The oocyte expands. B) Cartoon of ovariole structure.

One key question is the presence of individual germ stem cells which has been predicted by several authors^21,32,33^. Such germ stem cells, to follow the *Drosophila* model, would sit at the anterior end of the germarium, dividing as necessary. To search for the presence of these cells we focussed our scans on the anterior end of the germarium, attempting to determine if there is a population of single cells that may act as germ stem cells (Figure 2). In none of these scans could we find single cells that did not have the morphology of terminal filament cells, or were not attached, via a polyfusome, to a rosette of 8 cells. Germ stem cells have been postulated to appear in the terminal filament in honeybees^19^ and other bee species^18^. But without knowing which cells are germ-line and which somatic, it is hard to be clear where such a germ-cell population might reside.

**Figure 2:**
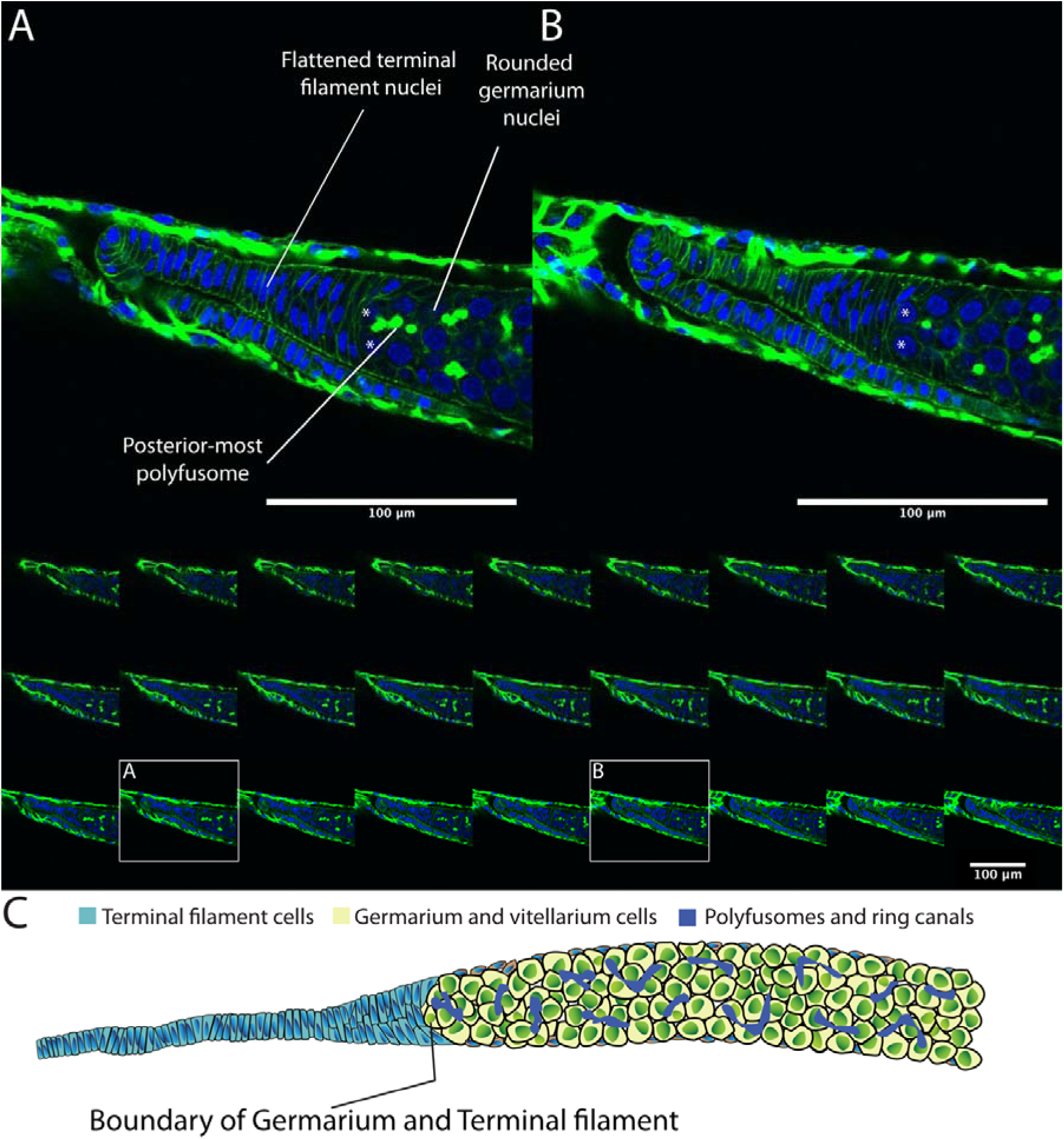
A) and B) single confocal sections across the boundary of the terminal filament and the germarium isolated from the montage of sections below. DAPI staining of nuclei (blue), and phalloidin staining (green) show the boundary between the flattened cells of the terminal filament and the rounded cells of the germarium. Germarium cells are joined by a polyfusome (staining with phalloidin). The two closest rounded cells to the terminal filament (marked with asterisks in A and B) are connected to surrounding cells by a polyfusome (most clearly shown in A). C) Cartoon of the boundary of the terminal filament and germarium drawn to show no germarium cells that are not attached to a polyfusome, even at the most anterior end of the germarium.

### Somatic and germ-line cells in the honeybee ovary

To determine which cells in the honeybee ovariole are germ-line, and which somatic, we used hybridisation reaction chain *in-situ* hybridisation^34,35^ to examine the expression of putative germ-line and somatic marker genes gleaned from the insect literature. Table 1 shows the genes we tested as potential markers and the evidence for their expression in each cell type.

RNA expression of *vasa* and *nanos* has long been established as germ-line markers in many animals (for examples see ^37,38,43,44,71–75^ including honeybees^22^. The expression of both these genes has been examined in honeybee embryos showing that germ-line cells can be identified in presumptive ovary tissue in late honeybee embryos^22^. RNA expression from these genes has also been examined in the ovary using histochemical methods, which did not produce a clear definition of the germ line in the germarium^22^. Using more sensitive HCR techniques^34,35^ (Figure 3) we found that both *vasa* and *nanos* expression are present in all the cells of the germarium, except the individual follicle cells on the outside of the germarium, and the cells of the terminal filament.

**Figure 3:**
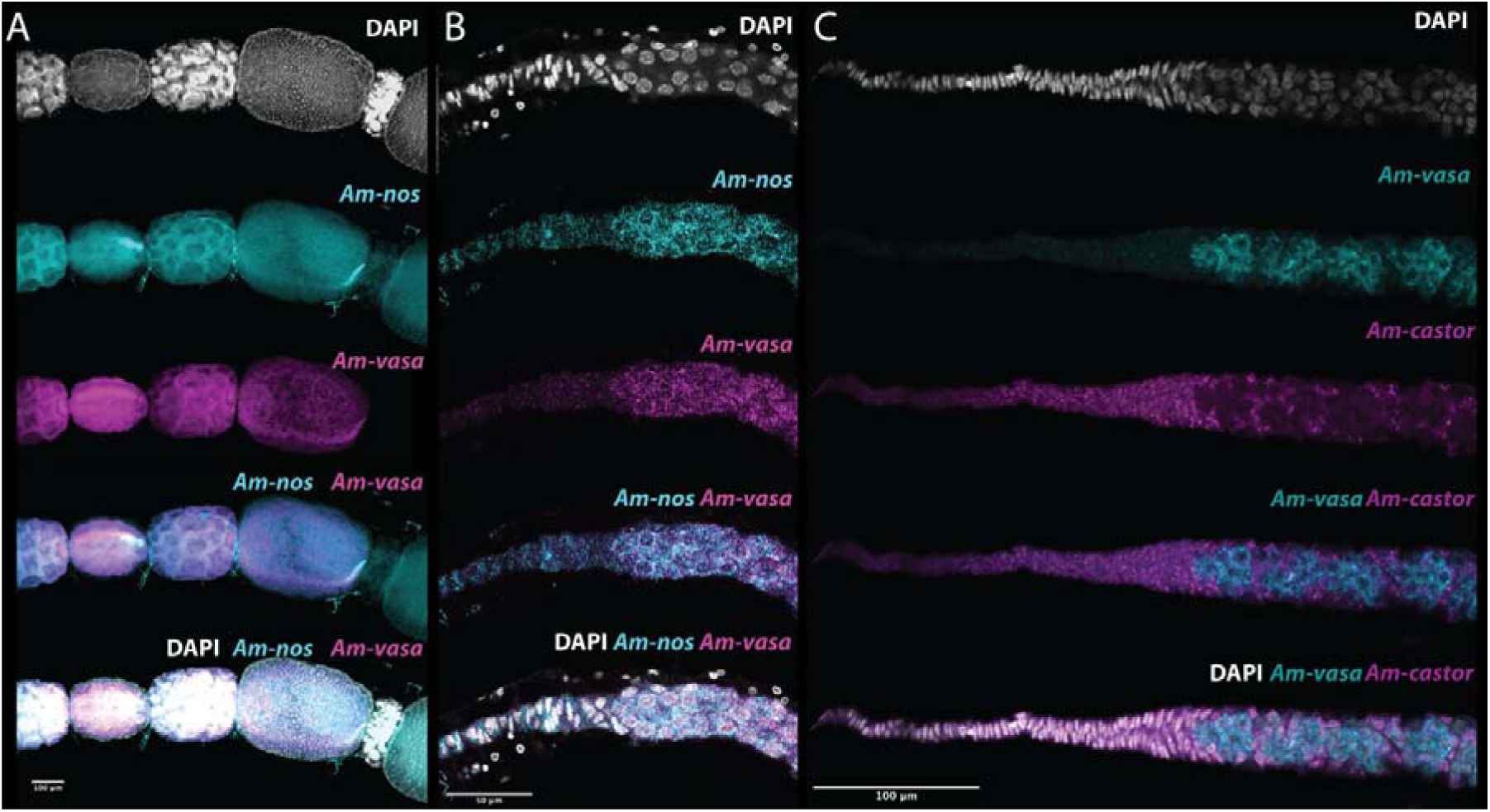
HCR in-situ hybridisation imaging of germ-line and somatic markers in honeybee queen ovarioles. A) Vitellarium region of an ovariole stained for *Am-nanos* (cyan), *Am-vasa* (magenta) and DAPI (grey). Expression of these conserved germline markers is consistent with previous publications^22^ with both germline markers expressed in nurse cells and the oocyte, with a concentration of RNA in a stripe down one side of the oocyte. There is no expression in follicle cells around the nurse-cell cluster or around the oocyte. B) Germarium and terminal filament region of the ovariole stained with *Am-nanos* (cyan), *Am-vasa* (magenta) and DAPI (grey). Both *Am-nos* and *Am-vasa* are expressed in the rounded germarium cells right up to the boundary with the terminal filament. No expression of either gene is present in the terminal filament. C) Germarium and terminal filament region of the ovariole stained with *Am-vasa* (cyan), *Am-castor* (magenta) and DAPI (grey). *Am-castor* expression is present in the terminal filament and follicle cells around the outside of the ovariole. Germ-line cells marked by *Am-vasa* are mutually exclusive to cells expressing *Am-castor*, implying *Am-castor* marks somatic cells in the ovary. As *Am-castor* is likely a somatic marker in the ovary^65,66^ this implies that no germ-line cells are present in the terminal filament.

In the vitellarium, all cells but follicle cells stain strongly for *Vasa* and *nanos* RNA (Figure 3), and, as the oocyte matures, the RNA of both genes comes to be located in a streak down one surface of the oocyte from the oocyte nucleus in the anterior, to the posterior of the oocyte (as previously reported^22^).

A few of our other potential markers showed similar gene expression in the germarium, including *dpp, ovo, eya* and *hh* (Supplemental table 1 and Supplemental Figure 2). *Ovo* in particular is expressed in very similar regions to *vasa* and *nanos*, making it a useful germ-line marker. Other potential markers, such as *Mad6*, also have expression in somatic follicles cells in *Apis*.

Somatic markers in the ovary have not been reported in insects outside of *Drosophila*, so we tested a panel of 6 markers, known to have somatic expression in Drosophila, to identify potential somatic cell markers. The honeybee ortholog of *Castor*, a somatic marker in *Drosophila*, is strongly expressed in the terminal filament, and in follicle cells in the ovary; an exact inverse of the expression of vasa and nanos (Figure 3). *Unzipped* RNA shows a similar pattern to *castor* in honeybees (Supplemental table 1 and Supplemental figure 1). In bees, *dpp, tj, hh* and *eya* all appear to have expression in some cells that are germ-line as well as somatic cells in the ovary (Supplemental table 1 and Supplemental figure 1), making them poor markers of the somatic cell population.

Identification of germ-line and somatic markers in the ovaries indicates that in honeybees, no germ cells are present in the terminal filament of the ovariole, as has been suggested by the previous authors^5^. The first germ-line cells of the ovary, as shown by the presence of *vasa* and *nanos* expression, and lack of *castor* expression, are 8 cell clusters, joined by a polyfusome, in the germarium, implying these are the stock of germ-line cells in the ovariole. We can identify no single vasa/nanos positive cells in the germarium, only 8-cell clusters joined by polyfusomes. In honeybee queen ovarioles, it appears that the germ line begins as clusters of 8 cells, physically joined by the polyfusome, located in the germarium.

That the stock of germ-line cells in the ovary are 8 cell clusters (rosettes in previous literature^4^), and that we can detect no single germ stem-cell in the ovary, led us to question how the high fecundity of queen bees is achieved. Are the ovaries of queen bees already populated with all the germ cells, as 8 cell clusters, that are needed after eclosion? Or are these 8 cell clusters capable of dividing? By examining markers of cell division, we aimed to rule out the presence of single dividing germline stem cells, and determine if the 8 cell clusters of germline cells can divide, and thus act as germ-line stem cells.

### Cell division in the honeybee ovariole

To begin to determine where cell division may be occurring in the germ-line of queen honeybee ovarioles we used immunohistochemistry against the mitosis marker phospho-histone Histone H3^76^ (pH3) to identify cells undergoing division. Cells undergoing division in the germarium of ovarioles appear in two places in each ovariole (Figure 4).

**Figure 4:**
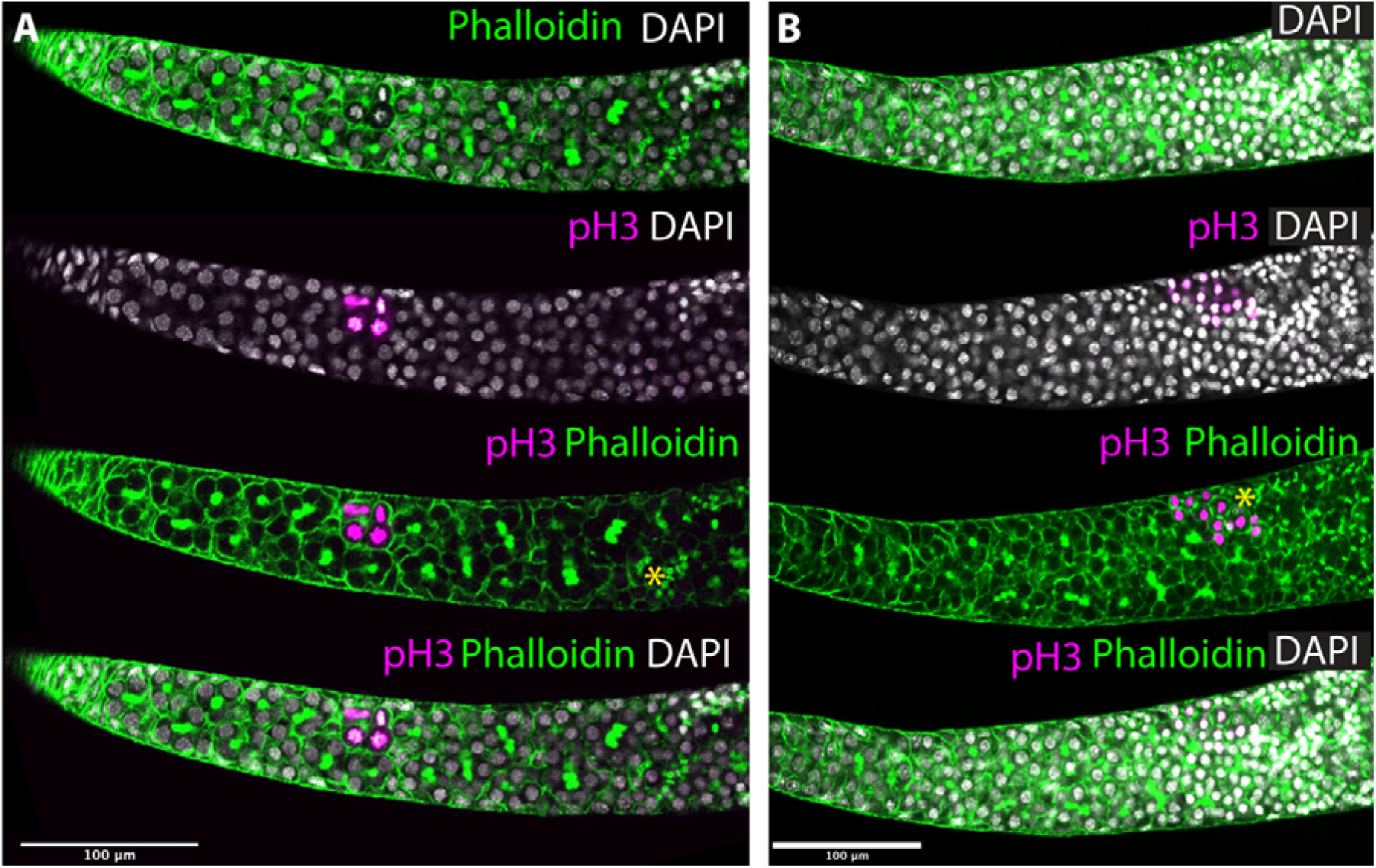
Immunohistochemistry for phospho-histone H3 in the germarium of honeybee ovarioles. A) Single confocal section of the anterior end of a germarium stained for phalloidin (green) and phospho-histone H3 (marking dividing cells (magenta)). In the anterior germarium one set of 8 cells (4 visible in this section) stains strongly with phospho-histone H3 (and can be seen to have metaphase chromosomes in DAPI). Asterisk marks the boundary between polyfusomes and ring canals. B) Single confocal section of the posterior end of a germarium stained for phalloidin (green) and phospho-histone H3 (magenta). Phospho-histone H3 positive cells can be seen at the boundary of the germarium, where polyfusomes give way to ring canals (boundary marked with an asterisk).

The first germarium region where mitosis is occurring is approximately halfway between the terminal filament and the posterior end of the germarium. Cells are often in mitosis in this area but always in clusters. These clusters, made up of 8 cells, appear to undergo mitosis in a synchronised way, producing two daughter 8-cell clusters from the original one. This data indicates that, despite the germ cells being linked by polyfusomes into 8-cell clusters, they are still able to divide, increasing the number of germ-cell clusters in each ovariole.

The second region is in the posterior germarium where polyfusomes are replaced by ring canals. As in this region, the oocyte is becoming specified, we believe this may be mitosis in the nurse cells as the oocyte undergoes the start of meiosis, with the oocyte remaining in metaphase I until oviposition^30^. We are yet to find a suitable meiosis marker in honeybees so are unable to confirm this.

We did not find, in any of our specimens, any signal from dividing single cells at the base of the terminal filament, as would be expected for a set of germ stem cells as in *Drosophila*. As we can find no single, non-terminal filament cells that stain with *vasa* or *nanos* and divide frequently, there is no evidence for a germ-line stem cell in the honeybee ovariole, apart from the 8-cell clusters in the germarium.

As our Ph3 staining indicates that the 8-cell clusters in the germarium can divide, this suggests that these are the germ-stem cells of the ovary, and key to the long-term fecundity of honeybees. It is remarkable that such clusters of cells, physically linked by a polyfusome, can divide. New markers for the polyfusome and more detailed scanning of these dividing cells are needed to address this question.

### Rate of germ-line division in honeybee germaria

The division of clusters of 8-cell germ-line clusters in the germarium of honeybees is caught only rarely by immunohistochemical staining for mitotic cells. To determine roughly how often these clusters divide we used EdU staining. EdU, a nucleotide analogue similar to BrdU, is incorporated into newly synthesised DNA in the S phase of cell division and remains detectable after cell division has occurred. This means by standardising the time between the treatment with EdU, and fixation and staining, we can get an estimate of how often 8-cell clusters divide in honeybee ovarioles. While it would be convenient to do this in tissue culture, we were concerned that ovarioles would react differently when removed from queen bees. To remove this concern we injected EdU into actively laying queens during spring and summer, allowed them to continue laying, and then euthanised queens and examined their ovaries 24 and 48 hours after injection (Figure 5).

**Figure 5:**
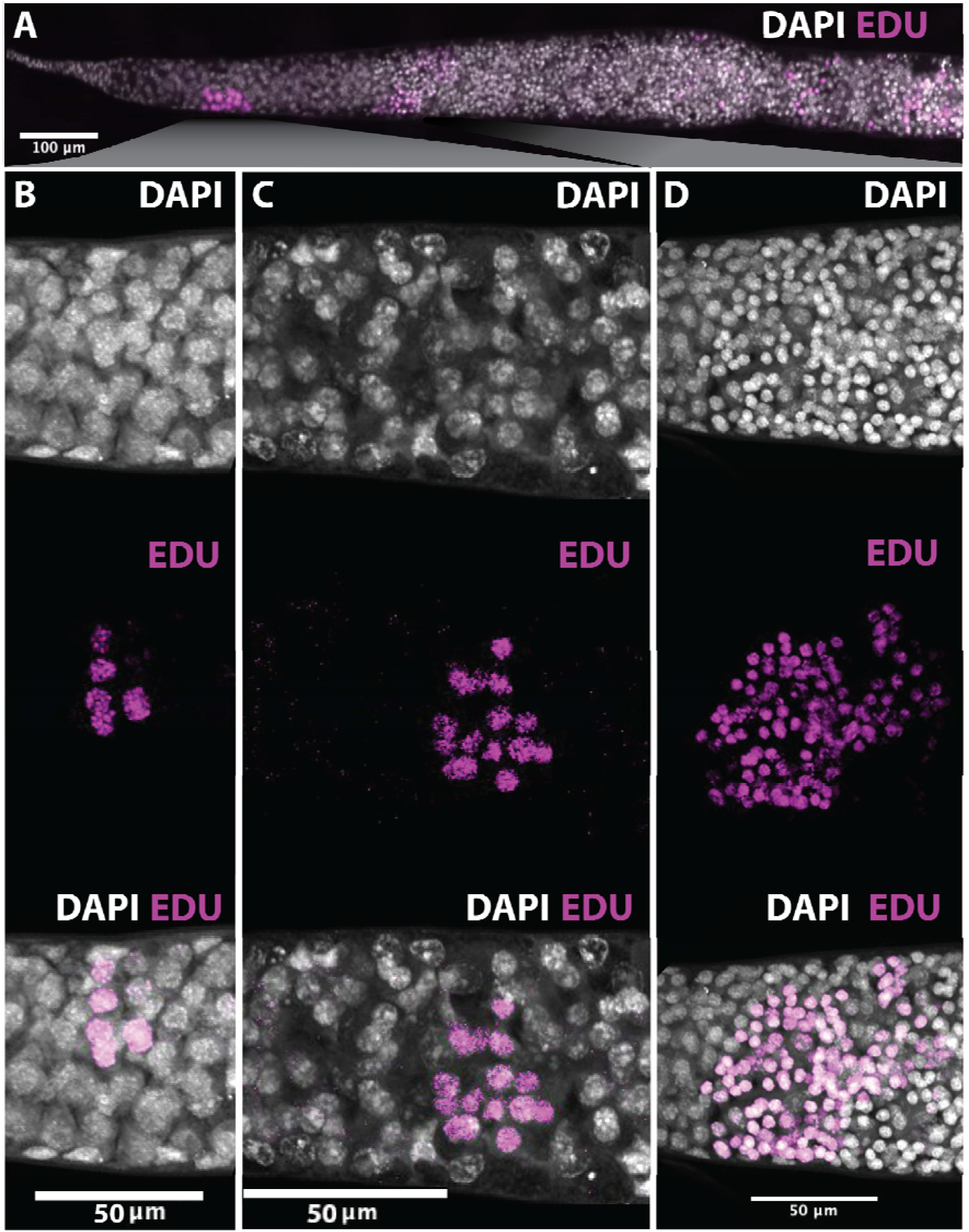
Detection of dividing cells using EdU after 24 hours of exposure. A) Terminal filament, germarium and anterior vitellarium of an ovariole from a queen bee injected with EdU, left to lay in a hive and fixed and stained after 24 hours. DAPI stained nuclei in grey and EdU, indicating DNA replication in the 24 hours (and thus cell division), in magenta. B) EdU staining in a cluster of 6-8 cells in the anterior of the germarium. C) A cluster of EdU marked 16 cells in the anterior of the germarium. D) a large cluster of EdU-marked cells at the boundary of the germarium and anterior vitellarium.

Examining ovarioles labelled and dissected after 24 hours indicates that, under these experimental conditions, a single cluster in the germarium divides once. Dissecting after 48 hours (Figure 6) provides evidence for two divisions in the germarium. The rate of division of the cells at the border of the germarium and vitellarium appears in the same range. In these conditions, and given the median number of ovarioles in each ovary (320^15^), and paired ovaries, this rate of division would reach that needed to produce the lower range of eggs^8–10^ expected from a laying queen in 24 hours.

**Figure 6:**
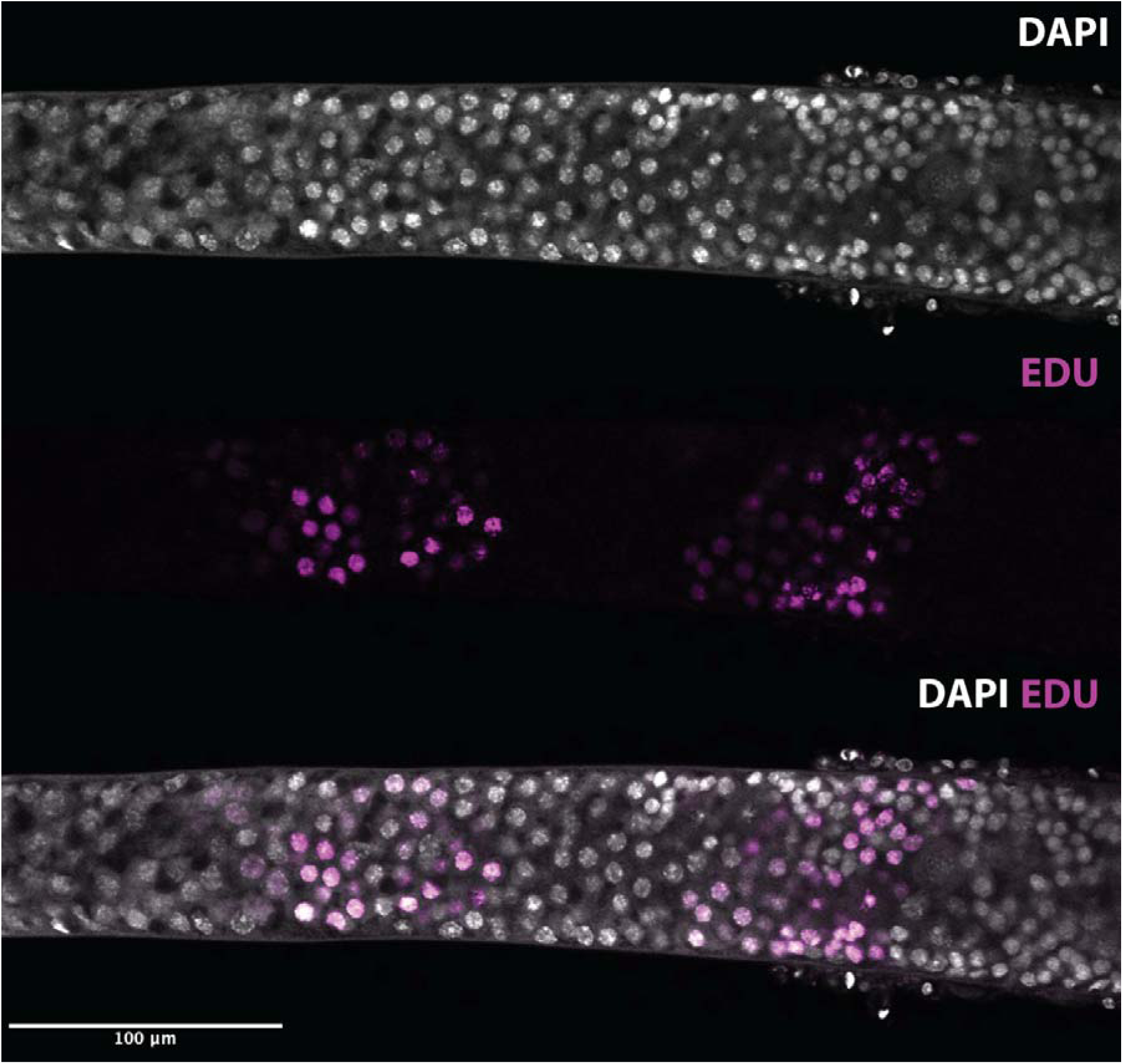
Detection of dividing cells using EdU after 48 hours of exposure. EdU staining in 2 clusters of cells in the anterior of the germarium and a large cluster of EdU marked cells at the boundary of the germarium and anterior vitellarium.

These are, by necessity, very rough figures as we cannot estimate how long it takes for the EdU injection to get to the ovary, how damaging the injection and handling is to queen reproduction (though all queens laid after injection), nor how much variation is added by local temperatures/weather conditions/hive placement etc. It is also clear that queens do not lay the maximum number of eggs possible **every** 24 hours, and indeed are reproductively inactive in cold weather. Our estimates are thus confounded by the significant environmental input into queen egg laying.

### Old queens do not have fewer germ-cell clusters than young queens

To determine if queen honeybees’ reproductive capacity is limited by the number of 8-cell clusters that form in their ovaries during larval development, we counted the number of polyfusomes in the germaria of young (3-10 month old) vs old (1-2-year-old) laying queens. Polyfusomes are easily detected by phalloidin staining, and so ovarioles from queens were stained with phalloidin, and confocal stacks were made. Polyfusomes were scored blind and independently by three experienced researchers. All the old queens in this experiment were used in normal commercial beekeeping operations and maintained outdoors near Dunedin, New Zealand. Old queens had slightly higher, but not statistically significant (p-value 0.159), number of polyfusomes (µ = 23.63, σ2 = 3.81) compared to old queens (µ = 22.49, σ2 = 4.46).

Our analysis shows no statistically significant difference in polyfusomes between young and old queens. This implies that the division of 8-cell clusters in the germarium occurs at a rate capable of replacing the stock of germ cells in queens over the long term. It seems likely that honeybee queens, at least over 1-2 years, do not run out of germ-stem cells in their ovary, and queen reproductive failure is more likely due to other effects (such as sperm depletion or damage).

## Discussion

Honeybee queens, like queens in many eusocial species, have remarkable reproductive capacity. This capacity is required to populate bee colonies over long periods and must be exquisitely responsive to the environment. This reproductive capacity has led to changes in the structure of the ovaries of honeybees, the most obvious being the large number of ovarioles in each of the two ovaries. The reproductive capacity of honeybees underpins natural and managed pollination around the world and the hive-produces industry.

Our data indicate that, unlike the model system *Drosophila*, there are no single germ-stem cells, as suggested by previous authors^21,32^, responsible for the production of oocytes located in the adult honeybee germarium. We cannot detect these cells in the ultrastructure of the ovary, nor do any such cells stain with germline markers, nor marked by phospho-histone H3, indicating active division, nor EdU, indicating division in at least 48 hours. Electron-microscopy^5^ and in-situ studies^22^ have suggested that germ-stem cells reside in the terminal filament of the honeybee ovary, but there are no cells in the terminal filament that stain for RNA from the germ-line markers *vasa* or *nanos*, and all these cells stain with our somatic marker *castor*. This arrangement is consistent with data from the polytrophic meroistic ovaries of other Hymenoptera^4^.

The germ-cell stock of the honeybee ovary appears then to be the collection of 8-cell clusters, linked by polyfusomes, that populate the germarium. These stain with the germ-line markers *vasa* or *nanos*, and are not somatic. This explains why bees have so many polyfusomes in each ovariole (average of 23) compared to *Drosophila*. In *Drosophila*, cystocytes with polyfusomes are a transient stage between germ-line stem cells and meiosis^17^. In honeybees, the 8-cell clusters, effectively the homologues of cystocyte clusters, are the germ-line stem population. These 8-cell clusters do not appear to form in the adult ovary but must be produced during larval and pupal development as the ovary forms, presumably from primordial germ cells which form late in embryogenesis^22^.

Given the germ-stem cells in the honeybee ovary are arranged in clusters, joined by polyfusomes, it would be unremarkable to suggest that these cannot divide and that the complement of the clusters in honeybee ovaries represents ALL the reproductive capacity of a queen bee. Our data shows this is not the case. The germ-stem-cell clusters can, and do, divide, and do so at a rate that is close to that required to maintain the reported reproductive capacity of queen bees. Examination of the germaria of old and young queens indicates that this rate of division is enough to maintain the reproductive potential of queens over their long lives. We find no sign of reproductive potential decline after 1 year of activity in queen bees, implying that, with good nutrition and a conducive environment, queen bees will remain highly reproductively active throughout their long lives.

How the conjoined clusters of germ-stem-cells achieve division with a polyfusome joining them is unknown, and will require more detailed observation of the division process with better markers for the polyfusome. Why division appears to be limited to a site about halfway along the germarium is also unknown, suggesting perhaps a permissive environment that exists at that site. We have found no evidence for differences in follicle cells, the intima membrane, or in RNA expression of the genes we have examined in this location that would suggest some change in the germ-stem-cell niche in this region.

Our identification of the germ-stem-cell niche in honeybees, and a large population of germ-stem-cell clusters in each ovariole, provides the potential for extracting, transplanting or even culturing these cells as a way to keep honeybee genetic stocks. Our finding that these cells have the potential to divide may provide the background knowledge to develop new reproductive technologies for these economically important insects.

## Author Contributions

GC: Experimental procedures, honeybee dissection, staining and injection, imaging and image analysis, manuscript drafting. JG: Polyfusome counting, manuscript drafting. JG: Statistics, manuscript drafting. PKD: Study conception, funding, image analysis, figure production, manuscript drafting, supervision.

## Acknowledgements

The authors would like to thank Dr Otto Hyink for supplying honeybees and supporting EdU injections. We thank B.P. Dearden for critical commentary and P.M. Dearden for critical reading of the manuscript.

