## Supplemental Figure 1 for "Germ-stem cells and oocyte production in the Honeybee Queen Ovary"

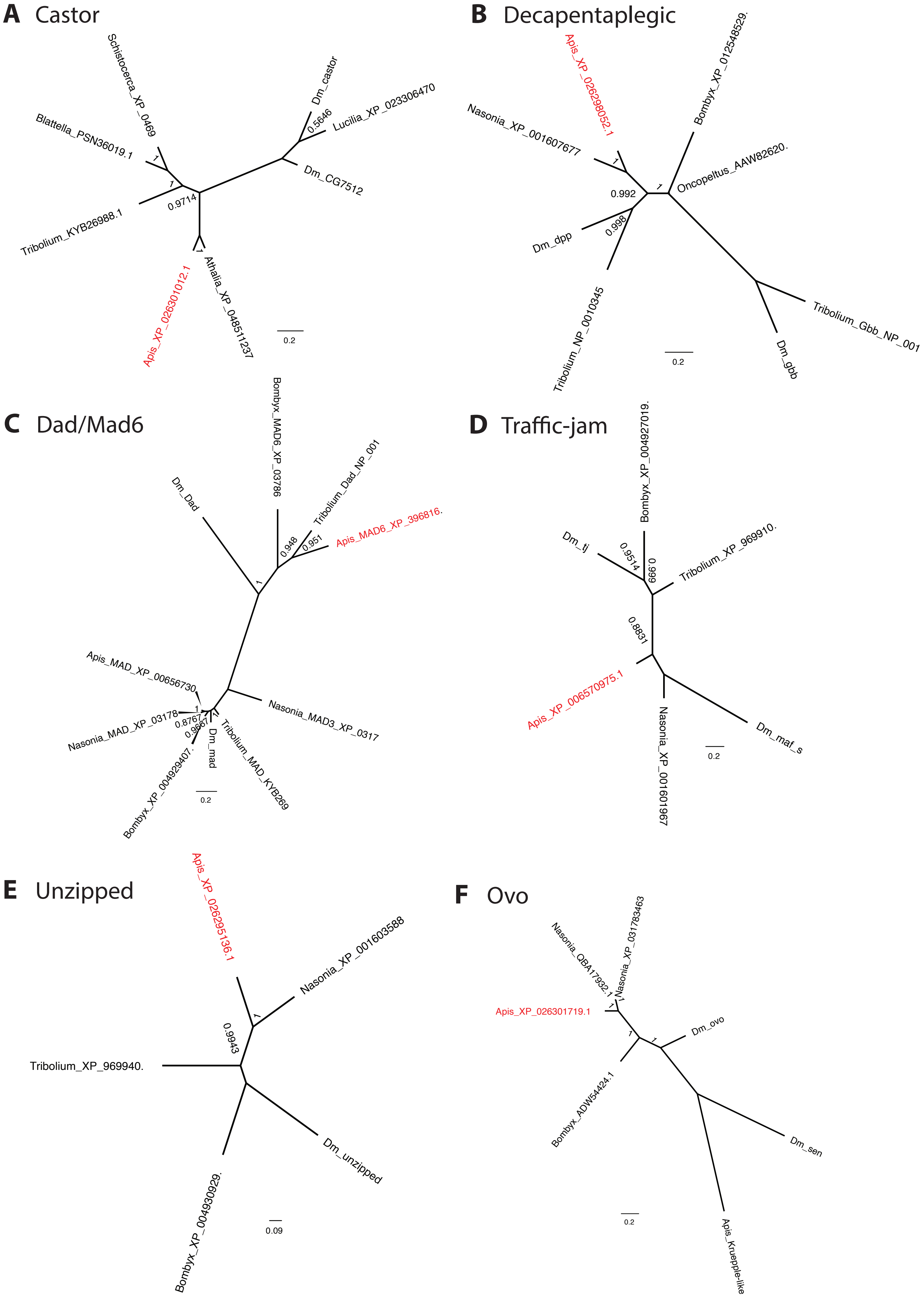

A Castor
Schistocerca_XP_0469
Blattella_PSN36019.1
Tribolium_KYB26988.1
Apis_XP_026301012.1
Athalia_XP_04851237
Dm_castor
Lucilia_XP_023306470
Dm_CG7512
0.5646
0.9714
1
0.2
B Decapentaplegic
Apis_XP_026298052.1
Nasonia_XP_001607677
Bombyx_XP_012548529.
Oncopeltus_AAW82620.
Dm_dpp
Tribolium_NP_0010345
Tribolium_Gbb_NP_001
Dm_gbb
0.992
0.998
1
0.2
C Dad/Mad6
Bombyx_MAD6_XP_03786
Dm_Dad
Tribolium_Dad_NP_001
Apis_MAD6_XP_396816.
Apis_MAD_XP_00656730
Nasonia_MAD_XP_03178
Nasonia_MAD3_XP_0317
Bombyx_XP_004929407.
Dm_mad
Tribolium_MAD_KYB269
0.948
0.951
0.8767
0.9667
1
0.2
D Traffic-jam
Bombyx_XP_004927019.
Dm_tj
Tribolium_XP_969910.
Apis_XP_006570975.1
Nasonia_XP_001601967
Dm_maf_s
0.9514
0.999
0.8831
0.2
E Unzipped
Apis_XP_026295136.1
Nasonia_XP_001603588
Tribolium_XP_969940.
Bombyx_XP_004930929.
Dm_unzipped
0.9943
1
0.09
F Ovo
Nasonia_QBA17932.1
Nasonia_XP_031783463
Apis_XP_026301719.1
Dm_ovo
Bombyx_ADW54424.1
Dm_sen
Apis_Kruepple-like
1
0.2

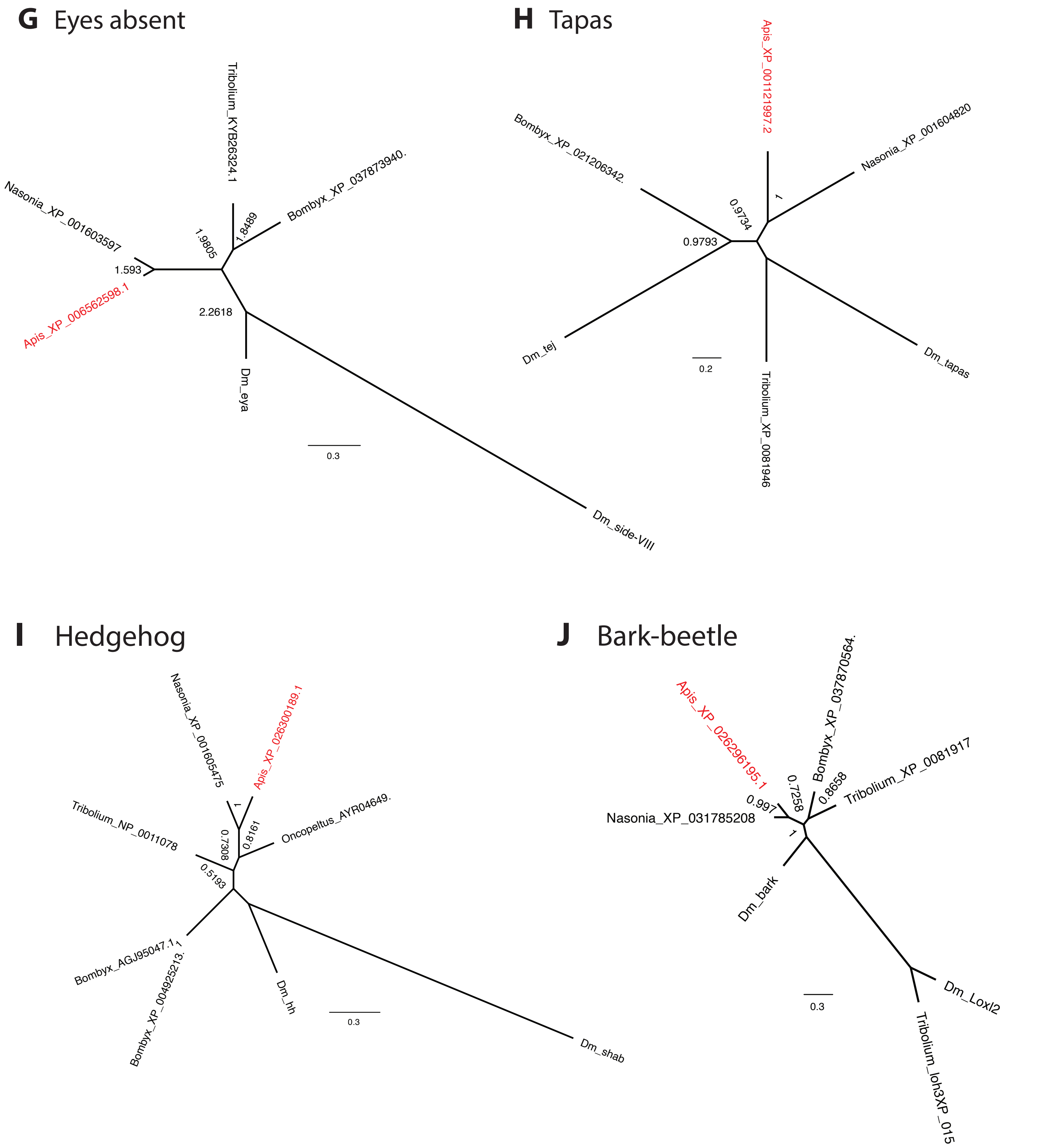


**Supplemental Figure 1:** Bayesian phylograms of ovary expressed genes. Phylogenies of Apis vasa and nanos are reported in REF 22 A) Honeybee *castor* (red) clusters with other *castor* orthologs against *Drosophila castor* and a closely related *Drosophila* protein *CG7512*. B) Honeybee *decapentaplegic* (red) clusters with other *dpp* orthogues against the related gbb . C) *Apis* Mad6 (red) clusters with *Drosophila* Dad against other MAD proteins. D) *Apis* Traffic jam (red) clusters with other Maf proteins. E) *Apis* unzipped (red) clusters with other unzipped proteins. F) *Apis* ovo (red) clusters with other ovo proteins and against closely related honeybee and *Drosophila* proteins. G) *Apis* eya (red) clusters with other eya proteins against *Drosophila* Side-VIII. H) *Apis* tapas (red) clusters with other tapas proteins against the closely related Tej protein. I) *Apis* hh (red) is closely related to other insect hh proteins to the exclusion of the related shab protein. J) Bark-beetle homologues, including Apis (red), clusetr to the exclusion of the related Loxl2 proteins.
