## Supplementary figures and images for "Germ-stem cells and oocyte production in the Honeybee Queen Ovary"

### Supplemental Figure 2

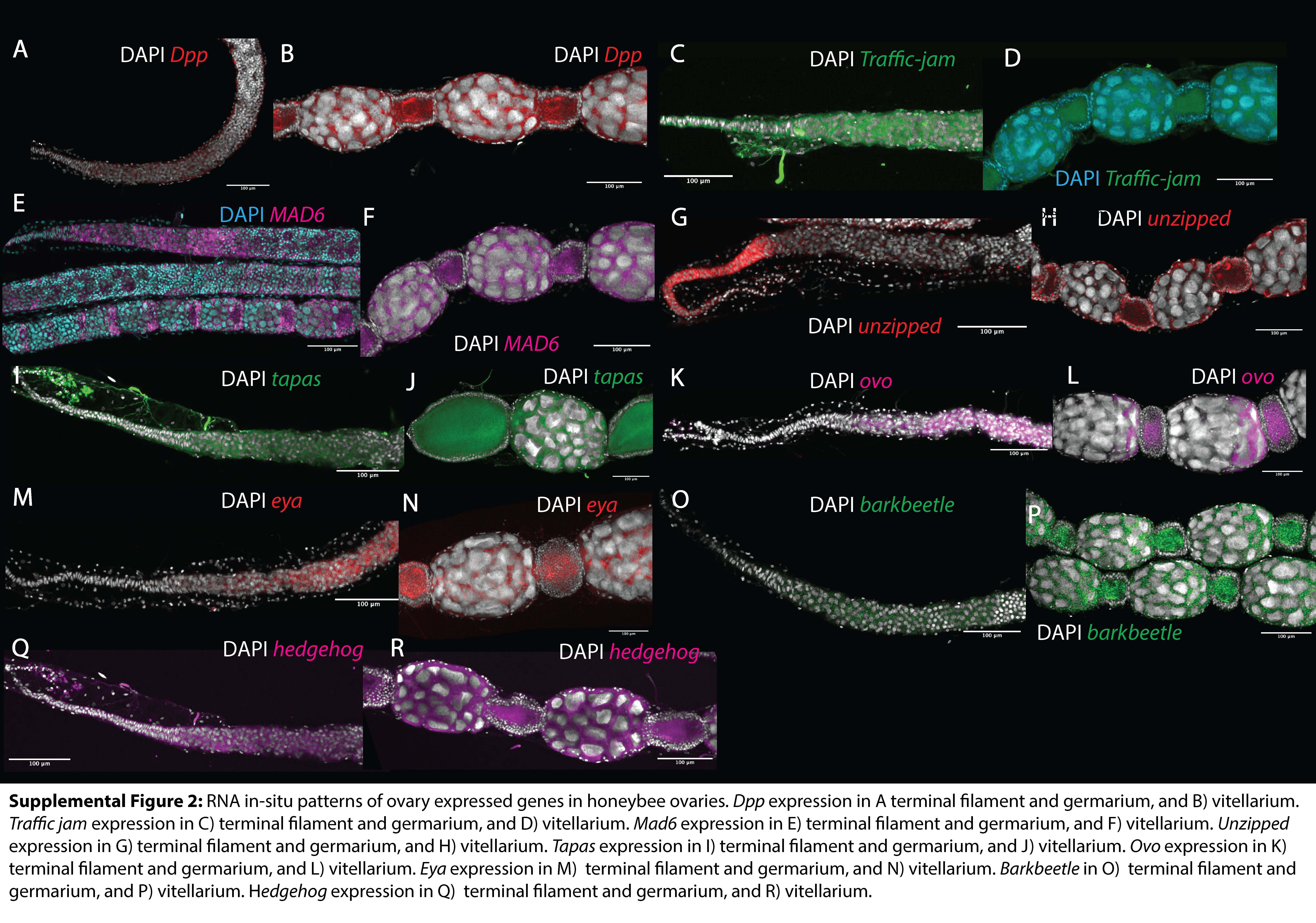
