## Supplemental Table 1 for "Germ-stem cells and oocyte production in the Honeybee Queen Ovary"

| **Gene homolog** | **Honeybee protein identifier** | **Queen ovary RNA expression pattern** | **Localised RNA in oocyte?** | **Somatic cells?** | **Germ-line cells?** | **Figure** |
| --- | --- | --- | --- | --- | --- | --- |
| *Vasa* | XP_006571765.2 | *Terminal filament:* no expression.  *Germarium*: germ-cell clusters. *Vitellarium*: nurse cells and oocytes | Yes, lateral stripe and posterior of oocyte | no | yes | 3 |
| *Nanos* | XP_026297805.1 | *Terminal filament*: no expression.  *Germarium:* in germ-cell clusters *Vitellarium:* nurse cells and oocytes | Yes, lateral stripe and posterior of oocyte | no | yes | 3 |
| *Castor* | XP_026301012.1 | *Terminal filament:* Strong expression  *Germarium and Vitellarium:* Follicle cells | No | yes | no | 3 |
| *dpp* | XP_026298052.1 | *Terminal filament:* no expression.  *Germarium:* In germ-cell clusters.  *Vitellarium:* some RNA in oocytes and nurse cells. | Yes, cortex of maturing oocytes and perinuclear | no | yes | S1 |
| *Traffic jam* | XP_006570975.1 | *Terminal filament:* weak expression.  *Germarium:* Expression in follicle and germ cell clusters.  *Vitellarium*: RNA in nurse cells and oocytes | No | yes | yes | S1 |
| *Mad6* | XP_396816.5 | *Terminal filament:* faint expression.  *Germarium:*  In follicle cells.  *Vitellarium:* in oocytes, particulate at the border of cysts, localised expression in late oocytes, Crescents of RNA in nurse cells and in follicle cells surrounding oocyte. | Yes, cortex of maturing oocytes and perinuclear | yes | yes | S1 |
| *unzipped* | XP_026295136.1 | *Terminal filament:* Strong expression.  *Germarium and Vitellarium:* follicle cells | no | yes | no | S1 |
| *Tapas* | XP_001121997.2 | *Germarium*: Nurse cells and oocytes | no | no | yes | S1 |
| *ovo* | XP_026301719.1 | *Terminal filament:* no expression.  *Germarium*: germ-cell clusters.  *Vitellarium:* Posterior nurse cells and oocytes | no | no | yes | S1 |
| *eya* | XP_006562598.1 | *Terminal filament:* no expression.  *Germarium*: Germ-cell clusters  *Vitellarium:* nurse cells and oocytes | no | no | yes | S1 |
| *Bark-beetle* | XP_026296195.1 | *Terminal filament:* no expression.  *Germarium:* no expression  *Vitellarium*: Posterior Nurse cells and oocytes | no | no | yes | S1 |
| *hedgehog* | XP_026300189 | *Terminal filament:* weak expression  *Germarium:* expression in germ clusters  *Vitellarium*: Some expression in follicle cells surrounding oocyte. Patchy expression in Nurse cells and oocyte | Yes, possibly cortical in oocyte | yes | yes | S1 |

**Supplemental Table 1:** Gene homologues used to identify somatic and germ-line cells in honeybee ovaries. Name column refers to the *Drosophila* gene name, except *MAD6*, which is named *Dad* in *Drosophila*. Accession numbers are for protein sequences. RNA sequences for these genes were used for the production of HCR probes.
